# A conserved regulation of cell expansion underlies notochord mechanics, spine morphogenesis, and endochondral bone lengthening

**DOI:** 10.1101/2024.08.12.607640

**Authors:** Brittney Voigt, Katherine Frazier, Donya Yazdi, Paul Gontarz, Bo Zhang, Diane S. Sepich, Lilianna Solnica-Krezel, Ryan S. Gray

## Abstract

Cell size is a key contributor to tissue morphogenesis^1^. As a notable example, growth plate hypertrophic chondrocytes use cellular biogenesis and disproportionate fluid uptake to expand 10-20 times in size to drive lengthening of endochondral bone^2,3^. Similarly, notochordal cells expand to one of the largest cell types in the developing embryo to drive axial extension^4–6^. In zebrafish, the notochord vacuolated cells undergo vacuole fusion to form a single large, fluid-filled vacuole that fills the cytoplasmic space and contributes to vacuolated cell expansion^7^. When this process goes awry, the notochord lacks sufficient hydrostatic pressure to support vertebral bone deposition resulting in adult spines with misshapen vertebral bones and scoliosis^8^. However, it remains unclear whether endochondral bone and the notochord share common genetic and cellular mechanisms for regulating cell and tissue expansion. Here, we demonstrate that the 5’-inositol phosphatase gene, *inppl1a*, regulates notochord expansion, spine morphogenesis, and endochondral bone lengthening in zebrafish. Furthermore, we show that *inppl1a* regulates notochord expansion independent of vacuole fusion, thereby genetically decoupling these processes. We demonstrate that *inppl1a*-dependent notochord expansion is essential to establish normal mechanical properties of the notochord to facilitate the development of a straight spine. Finally, we find that *inppl1a* is also important for endochondral bone lengthening in fish, as has been shown in the human *INPPL1*-related endochondral bone disorder, Opsismodysplasia^9^. Overall, this work reveals a conserved mechanism of cell size regulation that influences disparate tissues critical for skeletal development and short-stature disorders.

## RESULTS

Cell size is a key contributor to tissue morphogenesis^1^. Growth plate hypertrophic chondrocytes utilize cellular biogenesis and disproportionate fluid uptake to expand in size and lengthen endochondral bone ^2,3^. Defects of the growth plate can affect overall linear growth and contribute to skeletal disorders in humans, including both short– and tall-stature disorders^10^. Similarly, during vertebrate development, the osmotic swelling of notochord vacuolated cells (VCs), surrounded by an inelastic notochordal sheath, contributes to increased hydrostatic pressure and promotes elongation of the anterior-posterior axis^5^. This process establishes a stiff, rod-like axial tissue that provides leverage for locomotion and later acts as a template for spine development^11^. Cell size expansion contributes to tissue lengthening in both endochondral bone and the notochord; however, it remains unclear whether these two tissues share common genetic and cellular mechanisms for regulating this process.

In zebrafish, notochord VC expansion requires the fusion of small vacuoles to form a single fluid-filled vacuole that occupies the cytoplasmic space^7,8^. Defects in vacuolation or of the notochordal sheath render the notochord more deformable, leading to aberrant bending that can directly transition to vertebral malformations and scoliosis in adult animals^7,8,12^. Previous work has established that vacuole fusion is regulated by lysosome-related trafficking organelles and by V-ATPases, which facilitate fluid accumulation^7,8,13,14^. However, whether VCs expand by processes independent of vacuole fusion and whether these processes contribute to spine morphogenesis is unknown.

### Mutations in inppl1a cause thoracic scoliosis and vertebral malformations

In the continuation of our screen in zebrafish for mutations affecting adult spine development^15^, we identified a mutant in the 5’-inositol phosphatase encoding gene, *inppl1a*, that displayed severe thoracic scoliosis (**Figure S1**, see Methods and Materials). Homozygous *inppl1a*^*stl*445/*stl*445^ mutants (hereafter *inppl1a*^*stl*445^) were shorter in body length at 5 days post fertilization (dpf), and their body length remained shorter than wild-type controls throughout larval development (**Figure 1A-D**). During larval stages, *inppl1a*^*stl*445^ mutants displayed reduced viability, and surviving mutants developed notochord curvatures roughly above the swim bladder (**Figure 1E-H**). Adult *inppl1a*^*stl*445^ mutants were viable and displayed no obvious defects other than severe thoracic scoliosis in the dorsoventral plane (**Figure 1I-J**). Heterozygous *inppl1a*^*stl*445/+^ animals were phenotypically wild type at all stages of development. RNA sequencing analysis showed a significant loss of the *inppl1a* transcript in *inppl1a*^*stl*445^ mutants at 3 dpf, indicating that this allele is likely functioning as a null (**Figure S1J**).

**Figure 1.**
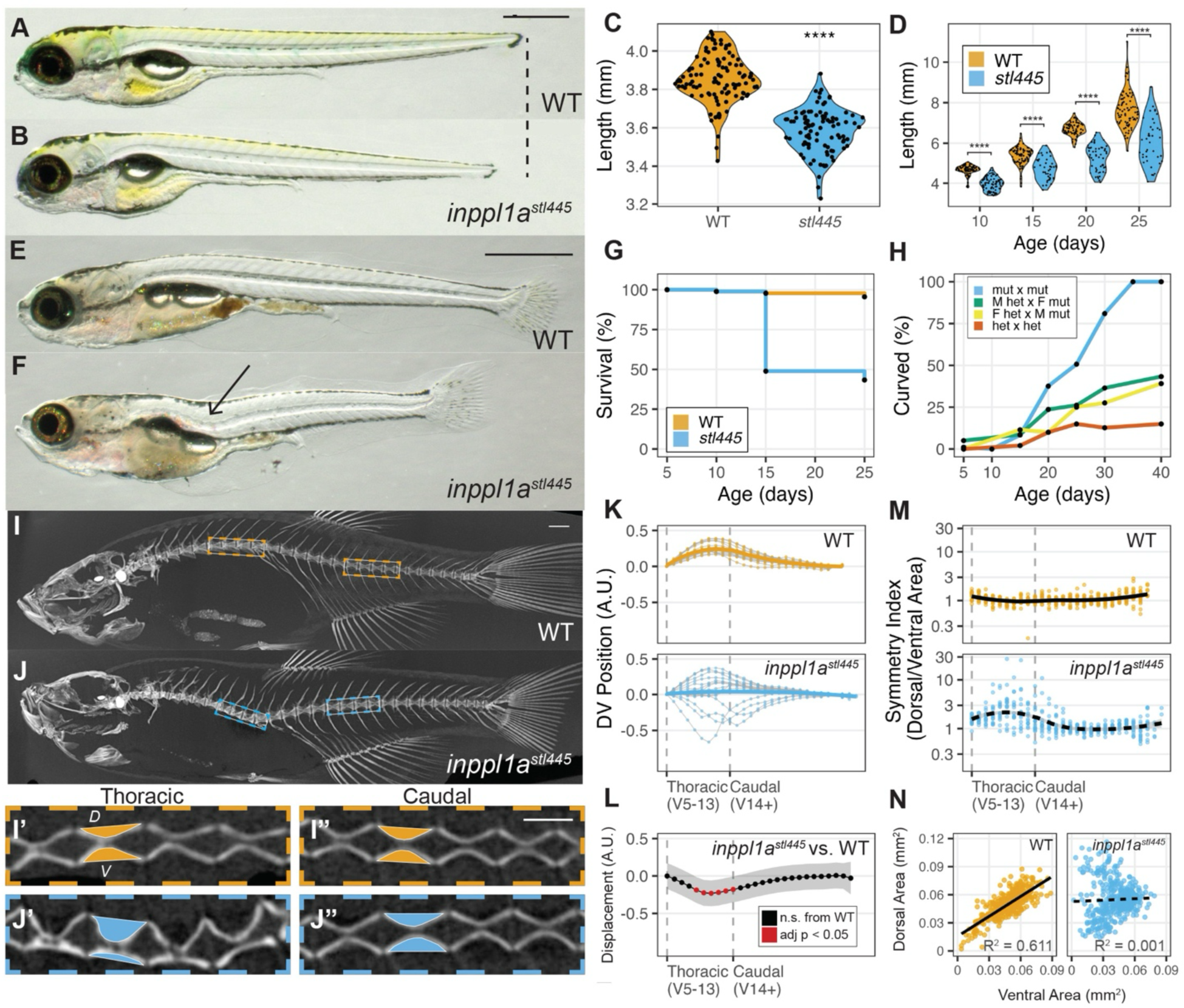
The inppl1a^stl445^ zebrafish mutant displays reduced body length, vertebral malformations, and thoracic scoliosis. (**A-B**) Representative wild type (A) and MZ *inppl1a*^*stl*445^ mutant (B) at 5 dpf. Scale bar = 0.5 mm. **(C)** Quantification of body length (mm) in wild type (n = 107) and MZ *inppl1a*^*stl*445^ mutants (n = 95) at 5 dpf (N = 3). **** indicates p-value < 0.0001. **(D)** Quantification of body length (mm) in wild type (n = 368) and MZ *inppl1a*^*stl*445^ mutants (n = 254) at 10, 15, 20, and 25 dpf. **** indicates p-value < 0.0001 by Tukey HSD test. **(E-F)** Representative wild-type (E) and MZ *inppl1a*^*stl*445^ mutant (F) juvenile fish with an arrow indicating the site of curvature. Scale bar = 1 mm. **(G)** Quantification of survival across larval development in wild type (n = 360, N = 3 independent crosses) and MZ *inppl1a*^*stl*445^ mutants (n = 360, N = 3). **(H)** Percentage of fish with curvatures from various crosses (homozygous mutant in-cross, n = 384, N = 5; male heterozygote crossed to female homozygous mutant, n = 238, N = 1; female heterozygote crossed to male homozygous mutant, n = 260, N = 2; heterozygous in-cross n = 200, N = 3) at 5, 10, 15, 20, 25, 30, and 40 dpf (± 2 days). **(I-J)** Representative maximum intensity projections of MicroCT imaging of adult wild type (I) and MZ *inppl1a*^*stl*445^ mutants (J). Scale bar = 1 mm. **(I’-J”)** Representative MicroCT midline projections of four thoracic (I’-J’) and caudal vertebrae (I”-J”) of fish pictured in I-J. The colored shapes represent the dorsal and ventral areas that are quantified in panel M-N. Scale bar = 0.5 mm. **(K)** Quantification of the normalized vertebral position (arbitrary units) on the dorsal-ventral axis in wild-type (n = 382 vertebrae, N = 16 fish) and MZ *inppl1a*^*stl*445^ mutant fish (n = 397 vertebrae, N = 15 fish) with the “loess” line of best fit in orange and blue. **(L)** Mean Displacement (arbitrary units) ± 95% Confidence Interval of MZ *inppl1a*^*stl*445^ mutant vertebrae compared to wild-type controls (as shown in panel K), colored by significance according to Tukey’s HSD Test. **(M)** Quantification of symmetry of each vertebra as measured by the dorsal area (mm^2^) divided by the ventral area (mm^2^) with loess lines of best fit for wild type (n = 382 vertebrae, N = 16 fish) and MZ *inppl1a*^*stl*445^ mutants (n = 397 vertebrae, N = 15 fish). Note the log scale on the y-axis. **(N)** Linear regression and correlation analysis of dorsal and ventral areas (mm^2^) of each vertebra in wild type and MZ *inppl1a*^*stl*445^ mutants (as shown in panel M).

We noted that the severity of the scoliosis phenotype in *inppl1a*^*stl*445^ mutants varied, so we developed a new method to provide a more granular quantification and statistical analysis of these spine defects using Micro-CT images. For this, we plotted the dorsoventral (DV) position of the centroid of each vertebra along the anterior-posterior axis of wild-type and *inppl1a*^*stl*445^ mutant fish (see details in Materials and Methods). Compared to the gentle dorsal-ward slope observed in wild-type zebrafish, our analysis of *inppl1a*^*stl*445^ mutants demonstrated a clear ventral-ward deviation of the spine that mostly affected the thoracic region (**Figure 1K**). Statistical analysis of these data using Tukey’s Honest Significant Difference (HSD) test found that vertebrae 9-13 were significantly displaced in the ventral direction in *inppl1a*^*stl*445^ mutants compared to wild-type controls (**Figure 1L**).

Some, but not all forms of scoliosis are associated with changes in the shape of individual vertebrae. Wild-type zebrafish displayed stereotypic hourglass-shaped vertebrae, with a high degree of symmetry between the dorsal and ventral sides (**Figure 1I’-I’’**). In contrast, we observed thoracic vertebral malformations in *inppl1a*^*stl*445^ mutants, as evidenced by increased cupping of the dorsal face and flattening of the ventral face of the vertebrae (**Figure 1J’-J”**). To understand how these vertebral malformations were associated with the presentation of scoliosis, we next developed a method to analyze the symmetry of individual zebrafish vertebrae. To do this, we defined a symmetry index as the midline area of the dorsal face of a vertebral body divided by the area of its ventral face (colored shapes in **Figure 1I”-J”**). By plotting this value for each vertebra, we observed a high degree of symmetry between dorsal and ventral faces along the length of the spine in wild-type fish (symmetry index near 1, **Figure 1M**). In contrast, *inppl1a*^*stl*445^ mutants displayed increased asymmetry with the greatest deviation within the thoracic region of the spine (**Figure 1M**). In addition, the correlation between dorsal and ventral areas for individual vertebrae was lost in *inppl1a*^*stl*445^ mutants (R^2^ = 0.001) compared to wild-type controls (R^2^ = 0.611, **Figure 1N**). Altogether, these results demonstrate that the localized incidence of vertebral asymmetry in *inppl1a*^*stl*445^ mutants (**Figure 1M**) was strongly correlated with the incidence of scoliosis in the same region of the spine (**Figure 1L**).

### inppl1a regulates notochord tissue and vacuolated cell expansion independent of vacuole fusion

During larval development, the notochord serves as a scaffold for vertebral bone deposition, which exerts concentric pressure on the notochord^8^. The mechanical stability of the notochord is directly dependent on processes regulating vacuole biogenesis and fusion, as well as extracellular matrix production by the notochord sheath^7,8,12^. Without sufficient notochordal hydrostatic pressure to push back against the forces of bone deposition, the notochord can collapse and lead to vertebral bone malformations and scoliosis in adult spines^8^.

Defects in notochord vacuole biogenesis and fusion can lead to shorter body lengths and smaller VCs containing fragmented vacuoles that become weak points during bone deposition and lead to vertebral malformations and scoliosis in adult zebrafish^7,8^. Similar to mutant zebrafish with disrupted vacuole fusion, *inppl1a*^*stl*445^ mutants displayed reduced body lengths, scoliosis, and vertebral malformations (**Figure 1**); however, when we visualized intracellular membranes using the vital dye, Cell Trace^7^, we found no evidence of defects in vacuole fragmentation or the appearance of notochord lesions at 3 or 5 dpf (**Figure S2A-D**). This result demonstrates that *inppl1a* is not essential for vacuole fusion in the notochord^7,8^.

To visualize if notochord and VC morphology was altered *inppl1a*^*stl*445^ mutants we used a fluorescent transgenic line Tg(*col8a1a:GFP-CAAX*^*pd*1152^) that specifically marks the plasma membrane of VCs^16^. We found that the notochord tissue in maternal zygotic (MZ) *inppl1a*^*stl*445^ mutants was shorter in length and smaller in diameter along the entire anterior-posterior axis compared to wild-type controls at 5 dpf (**Figure 2A-C, S3E-F**). We next quantified VC volumes using semi-automated 3D-cell segmentation (IMARIS) and found that the VC volumes were significantly reduced in *inppl1a*^*stl*445^ mutants compared to wild-type controls at 5 dpf (**Figure S2G-H**). These results demonstrate a role for *inppl1a* in VC size regulation that is independent of vacuole fusion and affects both axial and radial expansion of the notochord tissue.

**Figure 2.**
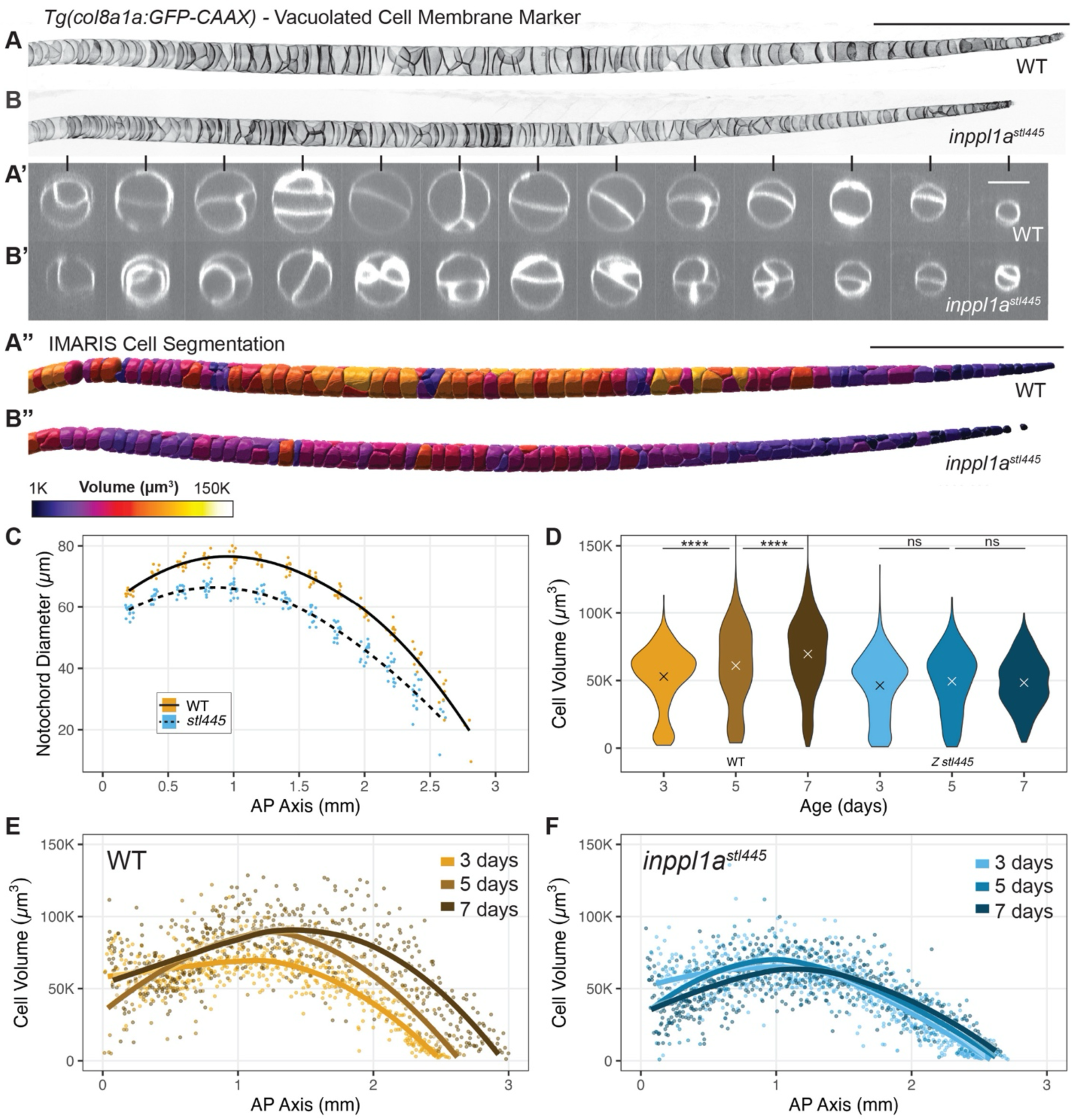
inppl1a regulates a phase of vacuolated cell swelling that is decoupled from vacuole fusion. **(A-B)** Representative max projections of confocal stacks of notochord vacuolated cell membranes (*Tg(col8a1a::GFP-Caax*)) in 5 dpf wild type (A) and MZ *inppl1a*^*stl*445^ mutants (B). Scale bar = 0.5 mm. **(A’-B’)** Orthogonal cross-sections of A-B at defined points along the anterior-posterior axis. Scale bar = 50 µm. **(A”-B”)** IMARIS Cell Segmentation of A-B colored by cell volume (µm^3^). Scale bar = 0.5 mm. **(C)** Quantification of notochord diameter (µm) in wild type (n = 10) and *inppl1a*^*stl*445^ mutants (n = 13) from orthogonal cross-sections as shown in A’-B’ with loess lines of best fit for each genotype. **(D)** Quantification of cell volume (µm^3^) at 3, 5, and 7 dpf in wild-type siblings (3 dpf: n = 494 cells, N = 4 fish; 5 dpf: n = 239 cells, N = 2 fish; 7 dpf: n = 538 cells, N = 4 fish) and zygotic *inppl1a*^*stl*445^ mutants (3 dpf: 655 = cells, N = 5 fish; 5 dpf: n = 239 cells, N = 2 fish; 7 dpf: n = 521 cells, N = 4 fish) from IMARIS cell segmentations. **** indicates adjusted p-value < 0.0001. The remaining adjusted p-values from Tukey’s HSD test are in Figure S2I. **(E-F)** Quantification of cell volume (µm^3^) (from panel D) along the anterior-posterior axis (mm) at 3, 5, and 7 dpf in wild type (E) and *inppl1a*^*stl*445^ mutants (F) with loess lines of best fit for each time point.

Wild-type notochord VCs generally complete the process of vacuole fusion by 2 dpf, and VC volumes exhibit a characteristic pattern along the anterior-posterior axis, cresting at the center of the notochord and then decreasing towards the tail^8^. Since vacuole fusion was not affected in *inppl1a*^*stl*445^ mutant embryos, we next sought to assess the dynamics of VC expansion after vacuole fusion. To do this, we imaged zygotic *inppl1a*^*stl*445^ mutants and wild-type siblings at 3 dpf, 5 dpf, and 7 dpf, and spatially mapped VC volumes along the anterior-posterior axis (**Figure 2D-F**).

Interestingly, the VC volumes of both wild type and *inppl1a*^*stl*445^ mutants exhibited a similar pattern along the anterior-posterior axis (**Figure 2E-F**). However, while the VCs undergo continuous expansion from 3-7 dpf in wild type larval zebrafish (**Figure 2D-E**), this process was arrested in *inppl1a*^*stl*445^ mutants (**Figure 2D & F** adjusted p-values in **Figure S2I**). In addition, our analysis also showed that *inppl1a*-dependent VC expansion contributes to the lengthening of the notochord from 3-7 dpf (**Figure 2E**), which is also absent in *inppl1a*^*stl*445^ mutants. Together these results demonstrate that *inpp1la* is essential for the phase of VC expansion that is independent of vacuole fusion and promotes radial expansion and axial lengthening of the notochord.

### inppl1a-dependent vacuolated cell swelling regulates notochord tissue mechanics during spine development

Because *inppl1a*^*stl*445^ mutants fail to expand VCs following vacuole fusion, we hypothesized that these mutant notochords may be more deformable and thus less resistant to the pressures of bone deposition. If true, this could contribute to notochord collapse during the onset of mineralization, which happens first in the thoracic region of the spine^17,18^, ultimately resulting in thoracic-localized vertebral bone malformations and scoliosis in adult mutants. To test this hypothesis, we reasoned that increasing the resistance to spontaneous movement and locomotion in *inppl1a*^*stl*445^ mutants during embryonic stages of notochord biogenesis may reveal mechanical instability of the notochord. To increase mechanical strain, we used a highly viscous 3% methylcellulose (MC) that has previously been used to exacerbate the effects of locomotion-induced notochord damage in mechanically sensitive backgrounds^16^. We found that wild-type embryos reared in 3% MC solution from 1-3 dpf (**Figure 3A**) displayed a significant decrease in VC volumes at 5 dpf, without disrupting the anteroposterior pattern of cell volumes (**Figure 3B-C, F-G**). Importantly, untreated *inppl1a*^*stl*445^ mutants displayed more reduced VC volumes when compared to 3% MC-treated wild types. However, 3% MC treatment of *inppl1a*^*stl*445^ mutants further enhanced the reduction of VC cell volumes and notochord length (**Figure 3D-F, H,** additional adjusted p-values in **Figure S3A**). This result demonstrates that exposure to increased mechanical resistance is sufficient to disrupt VC expansion and notochord lengthening, and further underscores the importance of *inppl1a* in this process.

**Figure 3.**
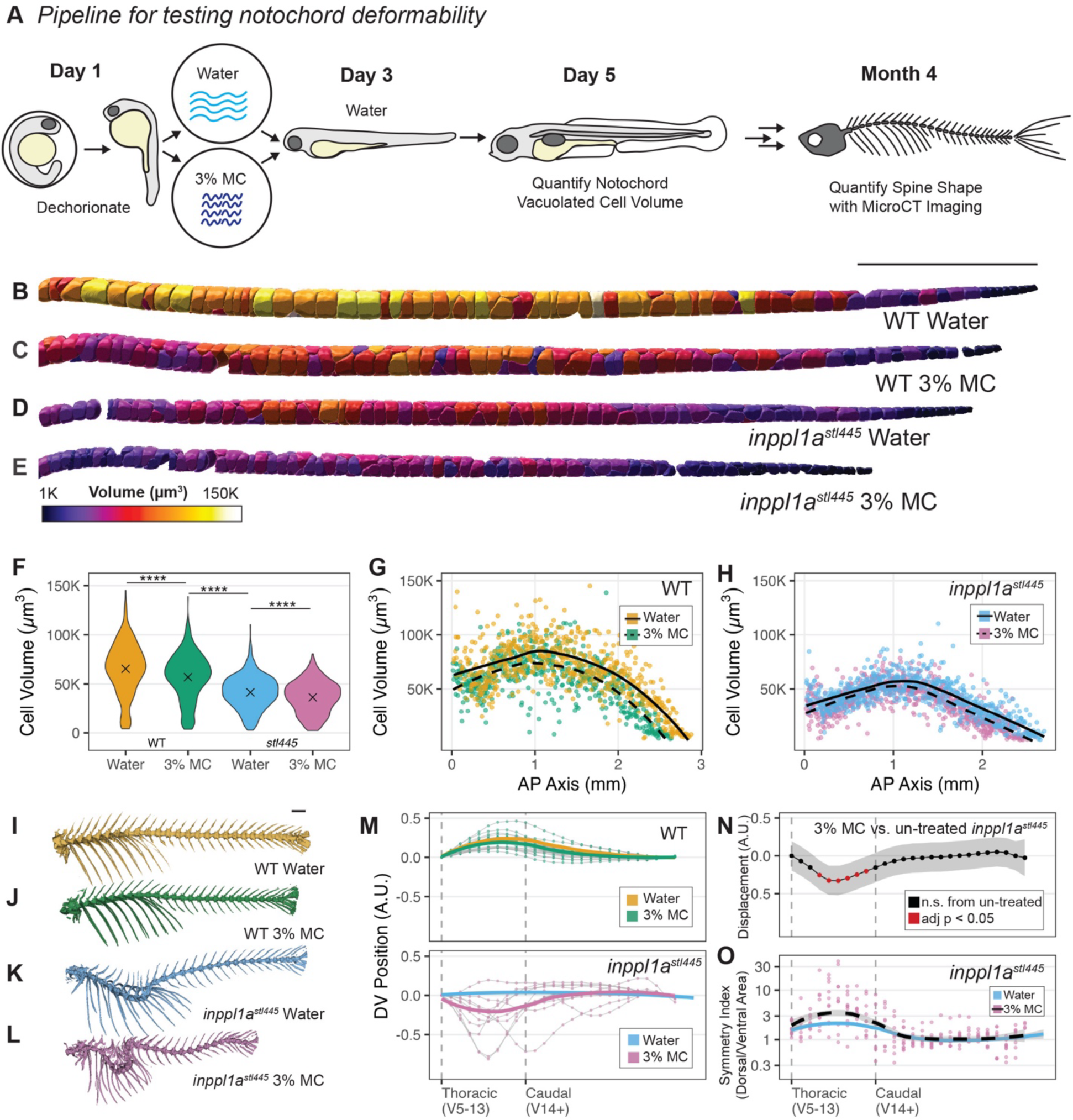
Disruption of post-vacuole fusion notochord expansion during embryonic development increases susceptibility to vertebral malformations and scoliosis in adult MZ inppl1a mutants. (**A**) Schematic showing the pipeline for testing notochord deformability using methylcellulose (3% MC) treatment with the analysis of notochord vacuolated cell volume at 5 dpf and spine shape at 4 mpf. **(B-E)** Representative IMARIS Cell Segmentation of notochord vacuolated cell-membrane marker *Tg(col8a1a:GFP-Caax)* of wild-type control (B), wild type methylcellulose-treated (C), MZ *inppl1a*^*stl*445^ mutant control (D), and MZ *inppl1a*^*stl*445^ mutant methylcellulose-treated(E) larvae at 5 dpf, colored by cell volume (µm^3^). Scale bar = 0.5 mm. **(F)** Quantification of cell volume (µm^3^) at 5 dpf of wild-type control (n = 601 cells, N = 5 fish), wild type methylcellulose-treated (n = 581 cells, N = 5 fish), MZ *inppl1a*^*stl*445^ mutant control (n = 646 cells, N = 5 fish), and MZ *inppl1a*^*stl*445^ mutant methylcellulose-treated fish (n = 755 cells, N = 6 fish). **** indicates adjusted p-value < 0.0001. The remaining adjusted p-values from Tukey’s HSD test are in Figure S3A. **(G-H)** Quantification of cell volume (µm^3^) (from panel F) along the anterior-posterior axis (mm) at 5 in wild type (G) and *inppl1a*^*stl*445^ mutants (H) with loess lines of best fit for each treatment. **(I-L)** Representative 3D renderings of MicroCT images of wild-type control (I), wild type methylcellulose-treated (J), MZ *inppl1a*^*stl*445^ mutant control (K), and MZ *inppl1a*^*stl*445^ mutant methylcellulose-treated (L) fish at 4 mpf. Scale bar = 1 mm. **(M)** Quantification of normalized vertebral position (arbitrary units) on the dorsal-ventral axis in methylcellulose-treated wild-type (n = 530 vertebrae, N = 21 fish) and MZ *inppl1a*^*stl*445^ mutant fish (n = 255 vertebrae, N = 10 fish) with the loess lines of best fit for each genotype in green and pink. Loess lines of best fit for control wild-type and MZ *inppl1a*^*stl*445^ mutant fish are superimposed from Figure 1 in orange and blue, respectively. **(N)** Mean Displacement (arbitrary units) ± 95% Confidence Interval of methylcellulose-treated MZ *inppl1a*^*stl*445^ mutant vertebrae compared to untreated MZ *inppl1a*^*stl*445^ mutant controls, colored by significance according to Tukey’s HSD Test. **(O)** Quantification of symmetry of each vertebra as measured by the dorsal area (µm^2^) divided by the ventral area (µm^2^) in MZ *inppl1a*^*stl*445^ mutants with loess lines of best fit in black. The Loess line of best fit for control MZ *inppl1a*^*stl*445^ mutant fish is superimposed from Figure 1 in blue. Note the log scale on the y-axis.

Given that defects in notochord biogenesis and expansion can contribute to vertebral malformations and scoliosis in adult fish^7,8,12^, we next wanted to test if increased mechanical resistance during embryonic notochord development would induce or enhance scoliosis in wild type or *inppl1a*^*stl*445^ mutants. We observed no obvious changes in spine morphogenesis in 3% MC-treated and untreated wild-type adult fish (**Figure 3I-J, M**). Statistical analysis did not reveal any evidence of differences of dorsoventral spine shape or vertebral malformations comparing 3% MC-treated and untreated wild-type animals (**Figure S3B-D**). These results suggest that some reduction of VC size may be tolerated as 3% MC-treated wild-type VCs were significantly larger than those of untreated *inppl1a*^*stl*445^ mutants (**Figure 3F**, additional adjusted p-values in **Figure S3A**). Alternatively, wild-type embryos may be able to expand VC size to restore normal notochord tissue mechanics after MC treatment.

In contrast, we found that exposure of *inppl1a*^*stl*445^ mutant embryos to 3% MC led to increased severity of scoliosis in these adults when compared to untreated mutants (**Figure 3K-M**). Statistical analysis of DV position revealed that thoracic vertebrae 8-12 were significantly more displaced in the ventral direction when comparing MC-treated to untreated *inppl1a*^*stl*445^ mutants (**Figure 3N**). Additionally, the 3% MC treatment led to more pronounced vertebral asymmetry in the thoracic vertebrae compared to untreated *inppl1a*^*stl*445^ mutant fish (**Figure 3O and S3D**). Therefore, by increasing mechanical strain during embryonic development, we have revealed that *inppl1a*-dependent regulation of VC expansion is essential to increase the hydrostatic pressure and mechanical stability of the notochord tissue during embryonic and larval development, which is ultimately required for the normal morphogenesis of the adult spine.

### inppl1a regulates endochondral bone lengthening in zebrafish

Homozygous mutations in the human *INPPL1* gene cause Opsismodysplasia (OPS), a rare endochondral bone disorder affecting growth plate expansion and skeletal development^9,19–22^. Therefore, we were interested to see if endochondral bones were affected in *inppl1a*^*stl*445^ mutant zebrafish. Unlike humans, however, zebrafish primarily use intramembranous bone development to build their skeletons, with only a few cranial endochondral bones^23,24^ and the appendicular hypural bones that connect the fin rays and appendages of the skeleton to the vertebral column^25^. Using Micro-CT imaging, we measured the lengths of the cranial intramembranous anterior parasphenoid (PS) bone^26,27^ and three endochondral hypural bones (the parhypural (PH), hypural 1 (H1), and hypural 2 (H2)) in the tail of wild-type and *inppl1a*^*stl*445^ mutant adults (**Figure 4A-D**). We found that both the PS and hypural bones were shorter in length in *inppl1a*^*stl*445^ mutants compared to age-matched wild-type controls (**Figure 4F-G**). To confirm that the decreased bone lengths were not just a reflection of an overall decrease in size of *inppl1a*^*stl*445^ mutants (as seen in **Figure 1**), we normalized the hypural lengths to the PS bone, which we found is correlated to measures of standard length in wild-type adults (**Figure 4E**). After normalization, we found that the ratio of the hypurals to the PS bone in individual fish was decreased in *inppl1a*^*stl*445^ mutants (**Figure 4H**). We repeated this analysis for the endochondral pharyngeal ceratohyal (CH) bones and found that these bones also exhibited a significant reduction in length (**Figure S4A-D**). Importantly, this analysis provides evidence for a conserved role for *inppl1a* in endochondral bone growth in zebrafish, as is already established in humans^20,28,29^ and mice^30,31^.

**Figure 4.**
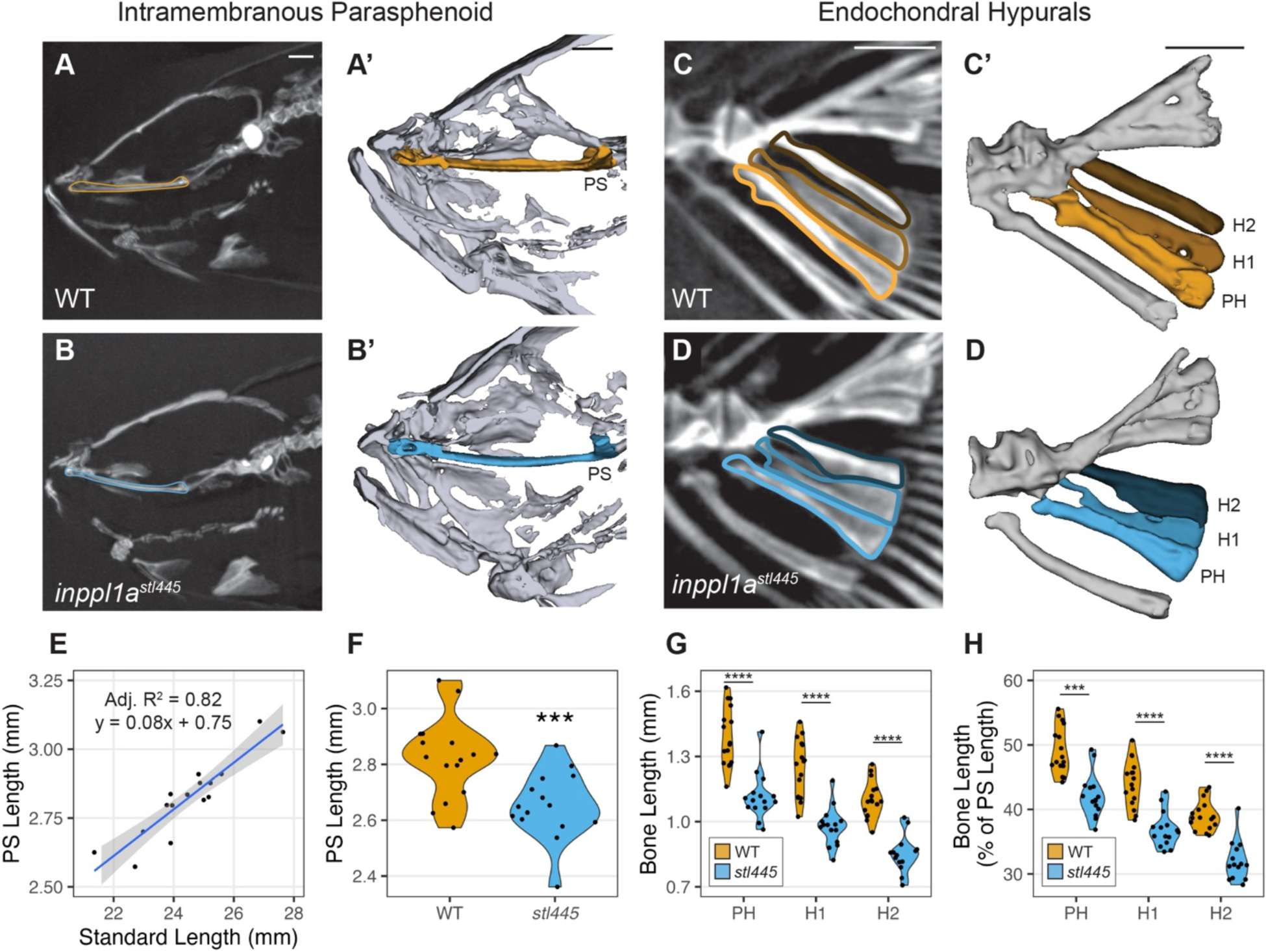
Endochondral bone lengthening is reduced in inppl1a^stl445^ mutants. **(A-B’)** Representative max projection of MicroCT data of the skull region surrounding the intramembranous anterior parasphenoid bone (outlined) in adult wild-type (A) and MZ *inppl1a*^*stl*445^ mutant fish, with corresponding 3D renderings (A’-B’). Scale bars = 0.5 mm. **(C-D’)** Representative max projections of MicroCT data of the tail region surrounding the endochondral hypural bones (outlined) in adult wild-type (A) and MZ *inppl1a*^*stl*445^ mutant fish, with corresponding 3D renderings (C’-D’). PH indicates parhypural, H1 indicates hypural 1, H2 indicates hypural 2. Scale bars = 0.5 mm. **(E)** Quantification of anterior parasphenoid (PS) length (mm) and standard length (mm) in adult wild-type fish, with corresponding linear regression line ± standard error (n = 16). **(F)** Quantification of anterior parasphenoid (PS) length (mm) in wild-type (n = 16) and MZ *inppl1a*^*stl*445^ mutant fish (n = 15). *** indicates p-value < 0.001. **(G)** Quantification of hypural bone lengths (mm) in wild-type (n = 16) and MZ *inppl1a*^*stl*445^ mutant adults (n = 15). PH indicates parhypural, H1 indicates hypural 1, H2 indicates hypural 2; **** indicates p-value < 0.0001. **(H)** Quantification of hypural bone lengths normalized as a percent of the anterior parasphenoid (PS) length in individual wild type (n = 16) and MZ *inppl1a*^*stl*445^ mutant adults (n = 15). PH indicates parhypural, H1 indicates hypural 1, H2 indicates hypural 2. *** indicates p-value < 0.001; **** indicates p-value < 0.0001.

## DISCUSSION

Inppl1a is a phosphoinositol-specific phosphatase that removes the 5-phosphate from phosphoinositide substrates in membranes (*e.g.,* PI_(3,4,5)_P_3_ and PI_(4,5)_P_2_)^32,33^. Previous *in vitro* experiments in a variety of cell culture models suggest that human *INPPL1* plays a role in the modulation of signaling cascades, cytoskeletal remodeling, and endocytosis^34–38^. AKT is a downstream effector of growth factor signaling and is a major regulator of cell proliferation, metabolism, and growth^39^. Similar to PTEN, Inppl1a is thought to oppose AKT signaling by depleting PI_(3,4,5)_P_3_, its primary activator^40^; however, other studies have indicated that the Inppl1a product, PI_(3,4)_P_2_, is required for full activation of AKT^41–43^. Additionally, decreased plasma membrane tension causes phase separation of PI_(3,4)_P_2_ which acts to recruit and inactivate TORC2 in budding yeast^44^. It is interesting to speculate that Inppl1-dependent generation of PI_(3,4)_P_2_ may help to regulate membrane tension and cell expansion in VC and chondrocytes.

In zebrafish, morpholino knockdown experiments suggest that *inppl1a* modulates fibroblast growth factor signaling important for embryonic dorsoventral patterning^45^. Against this notion, we did not observe any dorsalization or other embryonic patterning phenotypes either in our zygotic or maternal zygotic *inppl1a*^*stl*445^ or *inppl1a*^*ut*30^ mutant lines other than reduced body length and notochord VC swelling defects. However, general defects in growth factor signaling may contribute to the reduced growth rates and survival in homozygous *inppl1a*^*stl*445^ mutant clutches. While the exact mechanism of how Inppl1a regulates cell expansion remains unclear, we anticipate that additional effectors of Inppl1 may influence notochord development in zebrafish and endochondral growth plate development.

Cells can increase in size through true hypertrophy, which maintains overall cellular density via concomitant upregulation of macromolecules and organelles, or through osmotic swelling, where rapid cell size expansion is facilitated by disproportionate fluid intake^2^. Hypertrophic chondrocytes use both mechanisms of cell expansion^2^, while notochord VCs are thought to increase in size primarily through osmotic swelling^6^. As we see defects in both endochondral and notochordal tissues, our results suggest that Inppl1a is more important for cell swelling involving disproportionate fluid uptake. Furthermore, phosphoinositides such as the Inppl1a-substrate, PI_(4,5)_P_2_, can contribute to osmotic swelling through the regulation of ion channels and transporters that establish ion gradients across membranes^1,46^. However, since we observed that overall growth measures of *inppl1a* mutant animals were reduced compared to wild-type animals, our current study cannot rule out potential anabolic defects in *inppl1a*^*stl*445^ mutants that could also disrupt mechanisms underlying true hypertrophy.

Opsismodysplasia (OPS) is an autosomal-recessive endochondral bone disorder that causes congenital dwarfism, long-bone defects, craniofacial abnormalities, and in some cases, scoliosis^9,28^. Many of the OPS-related mutations in *INPPL1* are known to affect the catalytic domain, akin to the zebrafish *inppl1a* mutations reported here. Histological analysis of deceased OPS patients has revealed growth plate disorganization and shortening of the hypertrophic zones^20,28,29,47^. Similarly, mouse *INPPL1* mutants display shortened hypertrophic zones without changes in overall cell density, suggesting that the hypertrophic chondrocytes are smaller than their wild-type counterparts^30^. Inppl1-dependent processes underlie chondrocyte expansion in the mammalian growth plate and endochondral bone lengthening in zebrafish, defining a conserved role for cell size regulation across distinct cell types, tissues, and species. Building upon this knowledge, our study presents new evidence of a similar role for *inppl1a* in the notochord, such that *inppl1a* regulates VC size expansion that is independent of vacuole fusion and is also essential for normal lengthening and radial expansion of the notochord. Remarkable parallels exist between notochord and endochondral growth plate development: both involve convergent extension movements in progenitor cells to form linear arrangements of cells, and both utilize cell expansion to extend their respective tissues^48,49^. Altogether the conservation of Inppl1 for regulation of cell size during notochord maturation and endochondral bone formation suggests there may be additional shared genetic and cell biological processes that are essential for general cell expansion and tissue lengthening.

Overall, our findings highlight the role of *inppl1a* in the regulation of VC expansion in the zebrafish notochord, shedding light on additional mechanisms underlying notochord hydrostatic pressure establishment and spine development. We show that *inppl1a* regulates VC expansion that is independent of vacuole fusion and is also required for the development of a straight spine, as loss of *inppl1a* leads to adult thoracic scoliosis and vertebral malformations. We also demonstrate that increased mechanical stress during embryonic notochord development exacerbates the embryonic notochord VC size defect and severity of scoliosis and vertebral malformations in *inppl1a* mutant adults. Finally, we provide evidence that *inppl1a* plays a conserved role in the regulation of both notochord and endochondral bone lengthening. Further studies employing refined genetic approaches and characterization of Inppl1-effector proteins will be essential to fully understand this mechanism in the notochord and its relevance to chondrocyte hypertrophy.

## AUTHOR CONTRIBUTIONS

B.V. conducted all the experiments, performed all statistical analyses of *inppl1a* mutant phenotypes, generated all the figures, and wrote the original draft of the manuscript. K.F. and D.Y. assisted B.V. in crossing, genotyping, phenotyping, and performing various data analysis tasks. D.S.S. originally isolated the *stl445* zebrafish mutant in the L.S.K. lab. R.S.G. provided initial conceptualization for the project, supervised all experiments, and reviewed/edited the manuscript. B.V. (F31HD114419), L.S.K. (P01HD084387), and R.S.G. (R01AR072009) contributed to funding acquisition for this project.

## SUPPLEMENTAL INFORMATION

Figure S1. Mutations in *inppl1a* cause the *stl445 phenotype*.

Figure S2. Vacuole fragmentation and notochord vacuolated cell analysis in MZ *inppl1a*^*stl*445^ mutant larvae.

Figure S3. Notochord vacuolated cell and spine analysis in methylcellulose-treated wild type and MZ *inppl1a*^*stl*445^ zebrafish.

Figure S4. Bone length analysis of the endochondral ceratohyal bones in wild type and MZ *inppl1a*^*stl*445^ mutants.

Table S1. Allele frequency analysis of pooled *stl445* mutants and wild-type siblings.

Table S2. Differential gene expression analysis of *inppl1a*^*stl*445^ vs. wild type at 3dpf.

Document S1. Example code for vertebral position data import, manipulation, statistical analysis, and plotting in R.

## STAR METHODS

### Resource Availability

#### Lead contact

Further information and requests for reagents should be directed to the lead contact, Ryan S. Gray.

#### Materials availability

This study did not generate new unique reagents.

#### Data and code availability

RNA-seq data have been deposited at GEO and are publicly available as of the date of publication. Accession numbers are listed in the key resources table. Microscopy data reported in this paper will be shared by the lead contact upon request. The original code is available in this paper’s supplemental information (**Document S1**). Any additional information required to reanalyze the data reported in this paper is available from the lead contact upon request.

### Experimental Model and Subject Details

#### Zebrafish Maintenance and Care

All experiments were performed according to University of Texas at Austin IACUC standards. Wild-type AB strains were used unless otherwise stated. Embryos were raised at 28.5 °C in fish water (0.15% Instant Ocean in reverse osmosis water) and then transferred to standard system water at 5 dpf. Both male and female zebrafish were used for all adult assessments. As zebrafish do not exhibit sexual dimorphism until the early adult stages of development, sex is not a factor in any of the embryonic or larval assessments described in this manuscript.

### Method Details

#### Identification of stl445 scoliosis mutant

From the continuation of our screening in zebrafish for mutations affecting spine development^15^, we isolated the recessive *stl445* mutant that presented with scoliosis-like body curvature in the thoracic region. Through whole genome mapping and variant analysis (See *WGS / WES analysis*), a nonsense truncating mutation in the catalytic domain of the *inositol polyphosphate phosphatase like 1a* gene (*inppl1a;* c. T1261A; pY681*) on Chromosome 15 was identified as the cause of this phenotype (**Figure S1C-G; and Table S1**).

#### Skeletal preparation

Skeletal preparation was performed as in Wang *et al,* 2022^50^. Briefly, animals were fixed in 10% neutral buffered formalin and then incubated in 100% acetone overnight, and stained in Bone/Cartilage Stain (0.015% Alcian Blue, 0.005% Alizarin Red, 5% Acetic Acid, 59.5% Ethanol) at 37°C overnight. Tissue clearing was performed in 1% KOH for days to weeks depending on the size of the fish, followed by sequential glycerol embedding before imaging. For large fish, several images were captured and stitched together using the automated photomerge function in Adobe Photoshop or the pairwise stitching plugin in ImageJ/FIJI^51,52^.

#### WGS / WES analysis

Phenotypic, mutant zebrafish were pooled and submitted for sequencing. Non-phenotypic wild-type and heterozygous siblings were pooled together and submitted for sequencing. Raw reads were aligned to zebrafish genome GRCz10 using bwa mem (v0.7.12-r1034) with default parameters and were sorted and compressed into bam format using samtools (v1.6)^53^. Variants were called using bcftools (v1.9) functions mpileup, call, and filter. mpileup parameters “-q 20” and “-Q 20” were set to require alignments and base calls with 99% confidence to be used and filter parameters “-s LowQual –e ‘%QUAL<20’” were used to remove low quality variant calls. Variants were then annotated and filtered by an in-house pipeline. Briefly, variants occurring with the same allele frequency between phenotypic and wild-type samples were filtered from further analysis as were variants that were not called as being homozygous in the phenotypic sample. Variants in the mutant samples that were homozygous for the wild type allele were also excluded. Variants found in the dbSNP database (build v142) of known variants were also filtered out and excluded from further analysis. The wild type and mutant alleles at each variant site were tabulated, and the Fisher p-value was calculated for each variant site. These remaining variants were classified based on their genomic location as being noncoding site variants, coding site variants, or variants that may affect gene splicing using zebrafish Ensembl annotation build v83. For coding site variants, the amino acids of the wild-type allele and the mutant allele were determined from the Ensembl annotation. The p-values were plotted against genomic location, and a region of homozygosity in the genome with a cluster of small p-values was found. Variants occurring within this region of homozygosity were manually prioritized for nonsynonymous mutations in genes.

#### inppl1a^stl445^ genotyping method

The *inppl1a*^*stl*445^ mutation introduced an Hpy188III restriction site, enabling the molecular identification of heterozygous and homozygous carriers (**Figure S1F**). For genotyping, DNA was extracted from the tissue sample using the HotShot method^54^. Briefly, tissue samples were lysed in 50 mM NaOH and heated to 95 degrees C for 20 minutes before neutralization with a 1:4 volume of 1 M Tris-HCl (pH 8.0). Samples were either immediately used for PCR amplification or were stored at –20 degrees C until further processing. The *inppl1a*^*stl*445^ locus was amplified by PCR with Promega GoTaq Polymerase using the following primers: 5’-CTTCCTGTCATGTGACCTTCAG-3’ and 5’-GTTTGCATGTGTGTGTGTGTTC-3’ and the following settings: an initial denaturation at 95 degrees C for 3 minutes, followed by 34 cycles of denaturation at 95 degrees C for 30 seconds, annealing at 55 degrees C for 30 seconds, and extension at 72 degrees C for 30 seconds, followed by a final extension at 72 degrees C for 5 minutes. PCR samples were digested with Hpy188III (NEB R0622S) and screened on a 2% agarose gel to distinguish wildtype (double band), heterozygous (triple band), and homozygous *inppl1a*^*stl*445^ mutants (single band).

#### Generation of inppl1a^ut^^24^ allele

To verify the genotype-phenotype relationship between *inppl1a* and thoracic scoliosis, we generated a second mutation in the *inppl1a* gene via genome editing. CHOPCHOP^55^ was used to design a CRISPR guide RNA targeting exon 14 near the *inppl1a*^*stl*445^ locus (5’-GUCGCACCAUGAAGGAACGU-3’ with modifications to contain 2’-O’Methyl at the first three and last three bases and 2’ –phosphorothioate bonds at first three and last two bases from Synthego). One-cell stage embryos were injected with 1 nL of 5 µM equimolar sgRNA-Cas9 protein in 0.1 M KCl. Once the embryos reached sexual maturity, sperm was collected from F0 males to screen for INDEL founders^56^ using the same genotyping primers for the *inppl1a*^*stl*445^ allele (5’-CTTCCTGTCATGTGACCTTCAG-3’ and 5’-GTTTGCATGTGTGTGTGTGTTC-3’). PCR was done using GoTaq Green Master Mix (Promega #M7123) and amplicons were run on a 4% agarose gel. F0 founders were outcrossed, and F1 heterozygotes were in-crossed and screened for phenotypes. The new *inppl1a*^*ut*30^; c. C2001del8bp; p. N668F f.s.*24 allele also displayed recessive thoracic scoliosis and failed to complement the *inppl1a*^*stl*445^ allele (**Figure S1G-I**).

#### RNA Isolation, Sequencing, and Analysis

Total RNA was isolated from 3 dpf embryos using Trizol-reagent (Ambion by Life Technologies #15596026) and Direct-ZOL kit (Zymo #R2050). Library preparation, RNA sequencing, and bioinformatics analysis were performed by Novogene. Briefly. messenger RNA was purified from total RNA using poly-T oligo-attached magnetic beads. After fragmentation, the first strand cDNA was synthesized using random hexamer primers, followed by the second strand cDNA synthesis. The library was examined with Qubit and real-time PCR for quantification and bioanalyzer for size distribution detection. Quantified libraries were pooled and sequenced on Illumina platforms and paired-end reads were generated. Raw data in fastq format were first processed through in-house Perl scripts and clean reads were obtained. Reference genome and gene model annotation files were downloaded from the Ensemble genome website directly. The index of the reference genome was built using Hisat2 v2.0.5 and paired-end clean reads were aligned to the reference genome using Hisat2 v2.0.5. featureCounts v1.5.0-p3 was used to count the reads numbers mapped to each gene. FPKM of each gene was calculated based on the length of the gene and the reads count mapped to this gene. Differential expression analysis was performed using the DESeq2 R package (1.20.0). The resulting P-values were adjusted using Benjamini and Hochberg’s approach for controlling the false discovery rate. Genes with an adjusted P-value <=0.05 found by DESeq2 were assigned as differentially expressed (see Table S2). Gene Ontology (GO) enrichment analysis of differentially expressed genes was implemented by the clusterProfiler R package, in which gene length bias was corrected. GO terms with corrected p value less than 0.05 were considered significantly enriched in differentially expressed genes.

#### Larval Body Length Measurements and Curvature Assessments

Larval fish were anesthetized in 0.0016% Tricaine before imaging in 2% methylcellulose. Body length was measured in the lateral plane in ImageJ using the line tool, with anchors on the snout and posterior tip/flexion of the notochord depending on the developmental stage, as described in Parichy *et al.,* 2009^17^. To assess curvature, fish were collected from their tanks in a fine-mesh sieve and anesthetized in 0.0016% Tricaine. Fish were scored as either curved or straight from the lateral plane under a dissection microscope.

#### Micro-CT imaging and analysis

4-month-old fish were euthanized, fixed in 10% neutral buffered formalin, and mounted in 0.8% agarose for imaging. All samples were imaged on the Bruker Skyscan 1276 micro-computed tomography (Micro-CT) machine using the following X-ray parameters: 80 kV, 165 mA, 10.5 µm resolution, 0.4-degree step rotation (360-degree scan), 170 ms exposure time, and 2016×1344 image size. Raw data was reconstructed using the Bruker nRecon software (version 2.0) with the following parameters: 0.0-0.07 histogram, 0 beam hardening correction, 8 ring artifact reduction, and 3 smoothening. Reconstructed scans were rotated in Bruker Data Viewer software (v1.6.0.0) until they were optimally positioned for exporting sagittal sections jpeg datasets, which were imported into ImageJ as an Image Sequence for further analysis.

#### Vertebral Symmetry and Position Measurements

Beginning at the first thoracic vertebrae (V5), the dorsal and ventral vertebral face areas were measured from sagittal midline sections using the polygon tool in ImageJ. The vertebral position was measured at the midline using the multipoint tool in ImageJ. Position measurements were rotated and repositioned so that the Y coordinates for the first and 25th vertebrae were set to 0. To control for standard-length differences of individual fish^17^, we normalized the DV position to the length of the anterior parasphenoid bone, which we found is correlated to measures of standard length in wild-type fish (R^2^ = 0.836, **Figure 4E**), and its morphology was not altered in *inppl1a*^*stl*445^ mutants (**Figure 4B).** Specifically, each repositioned Y-coordinate was divided by the anterior parasphenoid length, which was measured from max projections of sagittal MicroCT images (**Figure 4A-B**). The symmetry and position data were imported into R for further analysis and plotting (See **Document S1**).

#### Other Skeletal Measurements

The hypural bones were measured in 3D Slicer^57^ (v5.6.2) using the fiduciary point tool on sagittal sections of MicroCT images. The standard length was measured from max projections of MicroCT images of whole fish.

#### 3D Skeletal Renderings

All renderings of MicroCT data were made in 3D slicer^57^ (v5.6.2). Folders of .bmp files were imported using SlicerMorph^58^ and segmentation of the skull, tail, and spines was performed using the Segment Editor feature at a threshold value of 80.

#### Confocal Microscopy and Analysis

Zebrafish embryos were placed in 0.003% phenylthiourea to block pigmentation at 1 dpf. Larval fish were anesthetized in 0.0016% Tricaine and mounted in 1% low melt agarose before imaging with the Nikon CSU-W1 Yokogawa Confocal Microscope. To obtain whole notochord images, a z-stack was taken throughout the entire depth and length of the notochord and automatically stitched together by Nikon software (NES Elements). All images were taken at 20X magnification with a 1-2 µm step size.

Notochord length was measured from maximum projections of these images, beginning at the anterior side of the swim bladder. Orthogonal sections were created in ImageJ, from which diameter measurements were taken at every 100 slices from the anterior side of the swim bladder. ND confocal stacks were uploaded into IMARIS (v10.0.1) for cell segmentation using the Cells and Surfaces tools. Briefly, all datasets were oriented in the same direction and a reference point was placed on the notochord just anterior to the swim bladder. The Cells module was used to segment the image using the following settings: Source channel 488 nm, cell smallest diameter of 15 µm (3 and 5 dpf) or 25 µm (7 dpf), membrane detail of 1.0 µm (3 and 5 dpf) or 2.5 µm (7 dpf), local contrast. Thresholding was manually set for each dataset, and once the cells were calculated, the results were manually edited to remove and/or fuse improperly segmented cells. The cell volumes were then exported to the Surfaces module for statistical measurements, including volume and distance from the origin reference frame. These datasets were exported and loaded into R for further analysis. Representative images were taken in IMARIS and were colored according to volume measurements.

#### Live BODIPY Staining and Imaging

At 1 dpf, 50 embryos per genotype were placed in 0.003% phenylthiourea to limit pigmentation. At 3 or 5 dpf, 10 individuals per genotype were transferred to Eppendorf tubes. The volumes were reduced to 250 µL, and 5 µL of BODIPY TR MED (Cell Trace) was added. Samples were incubated with open caps for 1 hour at 28.5 degrees Celsius. Before imaging, the samples were washed thrice in egg water, anesthetized in 0.0016% Tricaine, and mounted in 2% low-melt agarose. 20X magnification 50 µm confocal stacks were taken about the center of the notochord at the position dorsal to the yolk extension.

#### Methylcellulose experiment

Methylcellulose treatment was carried out as described previously^16^. Briefly, 1 dpf embryos were enzymatically dechorionated in Pronase for 5 minutes and randomly sorted into either egg water or 3% methylcellulose dissolved in egg water. Embryos were incubated at 28.5 degrees Celsius until 3 dpf, at which point they were transferred to egg water with gentle agitation to flush out the methylcellulose. Notochord imaging was performed as described above at 5 dpf (See *Confocal Microscopy and Analysis*). The experiment was repeated, and fish were allowed to grow up to 4 mpf at which point they were euthanized in excess tricaine for MicroCT imaging.

### Quantification and Statistical Analysis

All statistical analyses were performed in R (Version 4.3.2) using standard student t-tests or one-or two-way ANOVAs with Tukey Honest Significance Difference post hoc tests. Significance was defined as an adjusted p-value of less than 0.05 for all tests.

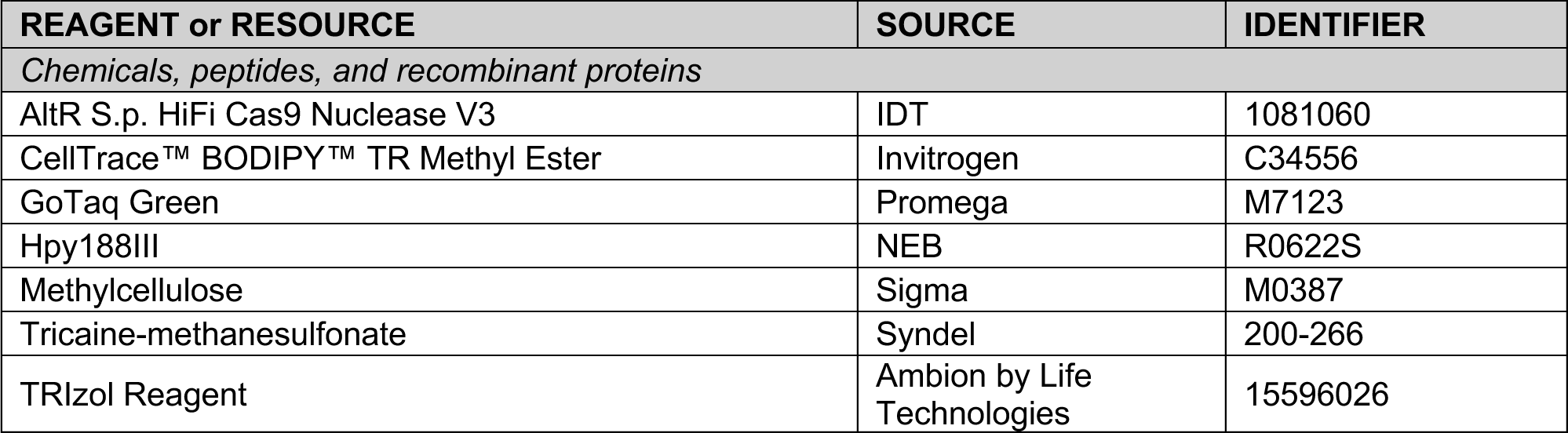
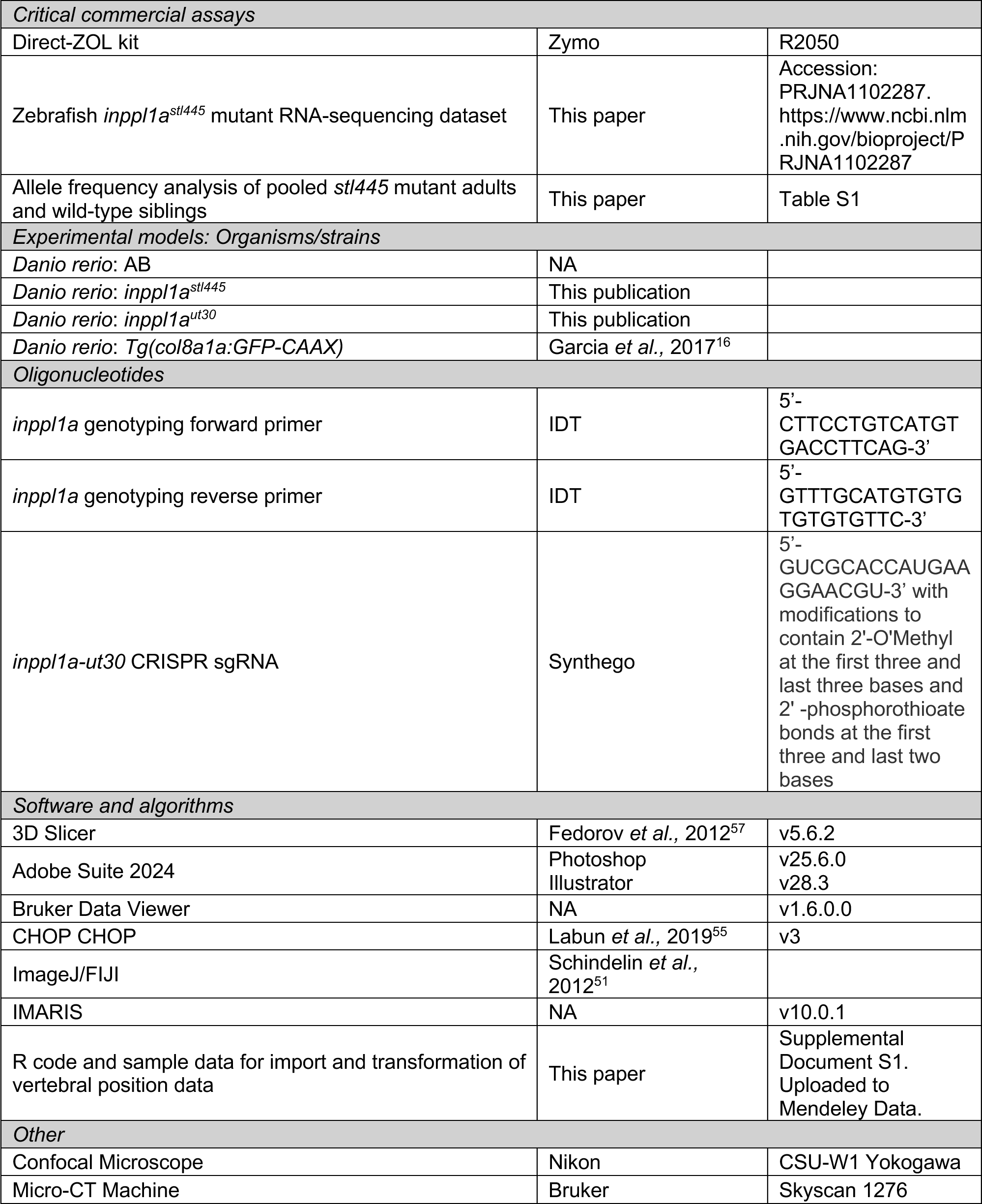
Key Resources Table.

## ACKNOWLEDGMENTS

We thank Michel Bagnat for sharing the *Tg(col8a1a:GFP-Caax)* vacuolated cell reporter line and members of the Gray lab for help with fish husbandry and helpful discussion. This work was supported by NIH under Awards R01AR072009 to R.S.G., P01HD084387 to L.S.K., and F31HD114419 to B.V.

---

**Figure S1.**
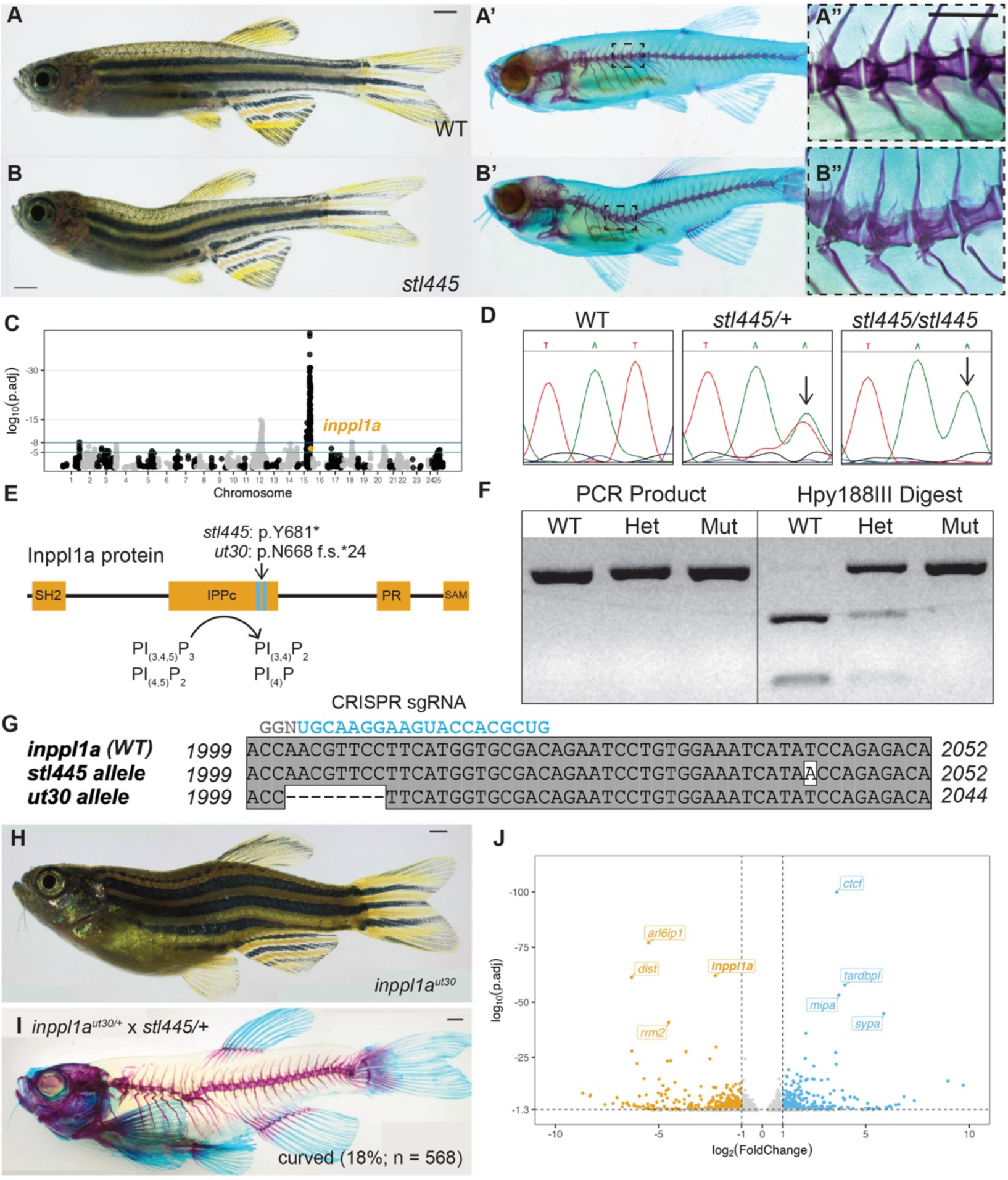
Mutations in inppl1a cause the stl445 phenotype. **(A-B)** Representative wild-type sibling (A) and zygotic *stl445* mutant (B) adults. Scale bar = 0.5 mm. **(A-B”)** Corresponding bone and cartilage stain (A’-B’), with inset showing four thoracic vertebrae (A”-B”, Scale bar = 0.5 mm). **(C)** Manhattan plot of SNP-linkage analysis from whole genome sequencing of *stl445* mutants (n = 20) versus wild-type control siblings (n = 20). Significance lines represent a log_10_(p-value) of –5 (green) and –8 (blue). The *inppl1a* SNP is highlighted in orange. **(D)** Representative Sanger sequencing results of wild type (n = 10), *stl445* heterozygous (n = 10), and *stl445* homozygous mutant (n = 20) fish at the *inppl1a^T2043A^* locus. **(E)** Domain architecture of the Inppl1a protein designating the SH2-homology containing domain, the inositol polyphosphatase catalytic (IPPc) domain, the proline-rich (PR) domain, and the sterile alpha motif (SAM) domain, as well as the approximate location of the ENU-generated *stl445* and CRISPR-generated *ut30* alleles. **(F)** Representative gel demonstrating the *stl445* genotyping method at the *inppl1a*^*stl*445^ locus. **(G)** DNA alignment of wild-type *inppl1a* with the ENU-generated *stl445* (T2043A) and the CRISPR-generated *ut30* (2001del.8bp) alleles, as well as the CRISPR sgRNA sequence in blue. **(H)** Representative homozygous zygotic *inppl1a*^*ut*30/*ut*30^ mutant fish. Scale bar = 1 mm. **(I)** Representative bone and cartilage stain of a curved trans-heterozygous fish from a complementation test between the *ut30* and stl445 alleles (18%; n = 568). Scale bar = 1 mm. **(J)** Volcano plot of differentially expressed RNA transcripts in 3 dpf MZ *inppl1a*^*stl*445^ mutants compared to wild-type controls (n = 40 per replicate, N = 3). Labeled genes represent the top four significant down-regulated (orange) and up-regulated (blue) genes in the dataset. Significance lines are designated at an adjusted p-value < 0.05 and a fold change greater than 2.

**Figure S2.**
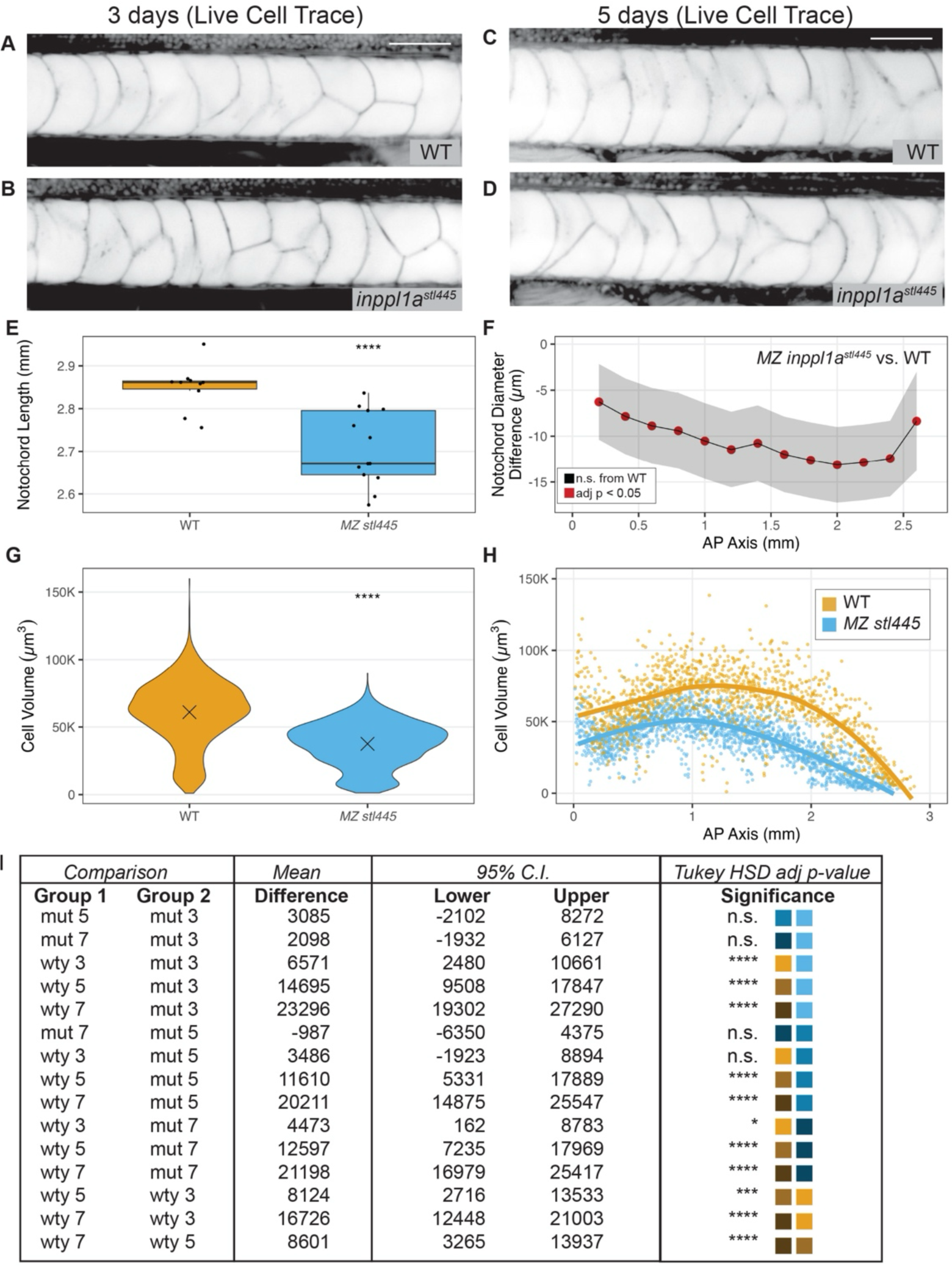
Vacuole fragmentation and notochord vacuolated cell analysis in MZ inppl1a^stl445^ mutant larvae. **(A-B)** Representative midline slices of 3 dpf wildtype (A, n = 4) and MZ *inppl1a*^*stl*445^ mutant (B, n = 8) larvae stained with Cell Trace BODIPY dye to mark intracellular membranes. Scale bar = 50 μm. **(C-D)** Representative midline slices of 5 dpf wild-type (C, n = 10) and MZ *inppl1a*^*stl*445^ mutant (D, n = 10) larvae stained with Cell Trace BODIPY dye to mark intracellular membranes. Scale bar = 50 μm. **(E)** Quantification of notochord length (mm) in wild type (n = 10) and MZ *inppl1a*^*stl*445^ mutants (n = 13) from confocal stacks as shown in A-B. **** indicates p-value < 0.0001. **(F)** Mean difference in notochord diameter (µm) ± 95% confidence interval from Tukey HSD test between wild type (n = 10) and MZ *inppl1a*^*stl*445^ mutants (n = 13) from orthogonal cross-sections as shown in Figure 2A’-B’. **(G)** Quantification of cell volume (µm^3^) at 5 dpf in wild type (n = 1253 cells, N = 10 fish) and MZ *inppl1a*^*stl*445^ mutants (n = 1855 cells, N = 13 fish) from IMARIS cell segmentations as shown in Figure 2A”-B”. **** indicates adjusted p-value < 0.0001. **(H)** Quantification of cell volume (µm^3^) (from panel G) along the anterior-posterior axis (mm) at 5 dpf in wild type and maternal zygotic (MZ) *inppl1a*^*stl*445^ mutants with loess lines of best fit for each genotype. **(I)** Results of Tukey HSD test comparing notochord VC volumes in various groups as shown in Figure 2D. Colored squares match the color palette of Figure 2D. *, ***, and **** indicate adjusted p-values < 0.05, 0.001, and 0.0001, respectively.

**Figure S3.**
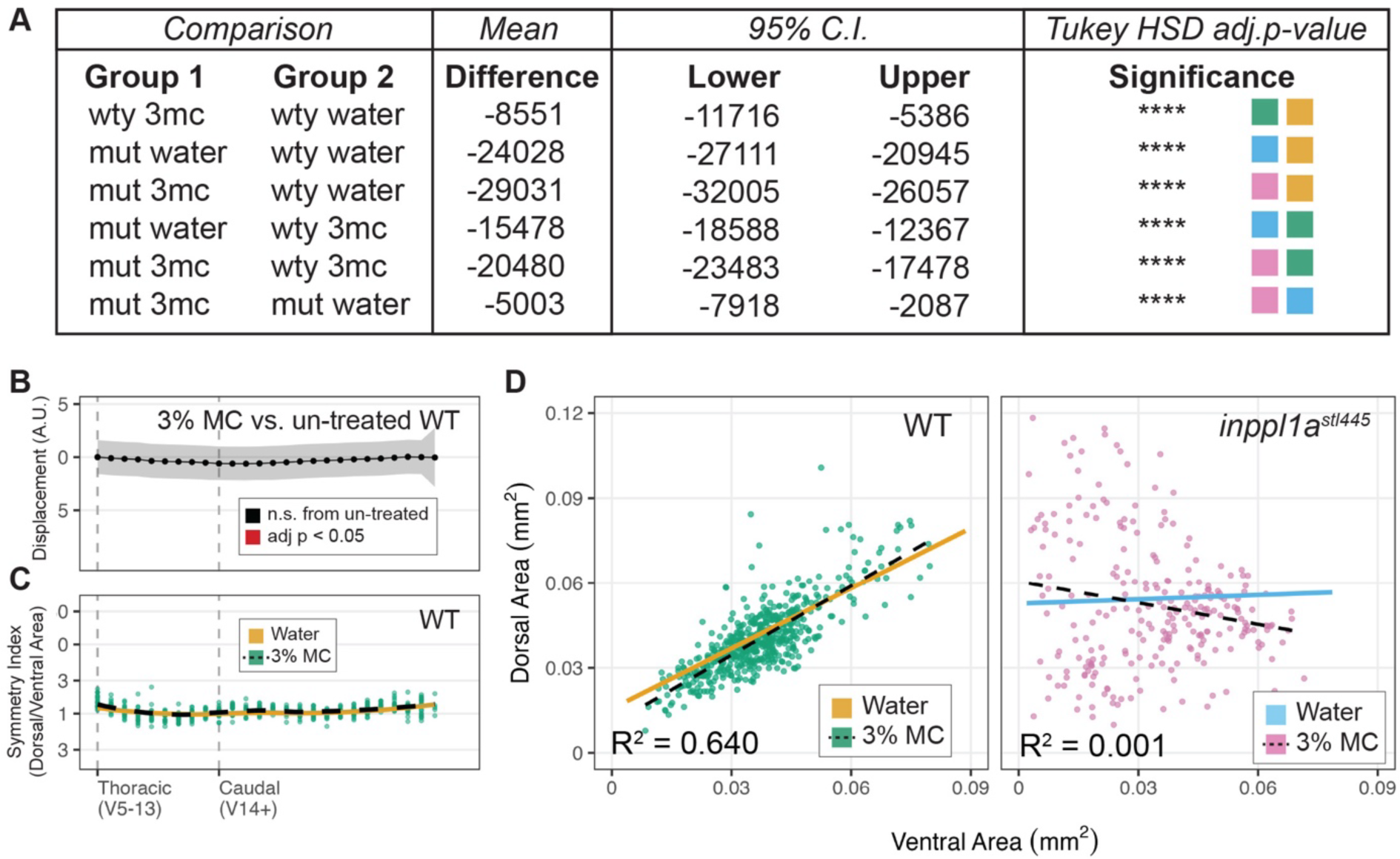
Notochord vacuolated cell and spine analysis in methylcellulose-treated wild type and MZ inppl1a^stl445^ zebrafish. **(A)** Results of Tukey HSD test comparing notochord VC volumes in various groups as shown in Figure 3F. Colored squares match the color palette of Figure 3F. **** indicates adjusted p-value < 0.0001. **(B)** Mean Displacement (arbitrary units) ± 95% Confidence Interval of methylcellulose-treated wild-type vertebrae compared to untreated wild-type controls, colored by significance according to Tukey’s HSD Test. Note that none of the vertebrae are significantly displaced. **(C)** Quantification of symmetry (arbitrary units) of each vertebra as measured by the dorsal area (µm^2^) divided by the ventral area (µm^2^) of methylcellulose-treated wild-type fish, with dashed loess line of best-fit fin black. The Loess line of best fit for wild-type control is superimposed from Figure 1 in orange. Note the log scale on the y-axis. **(D)** Quantification of dorsal and ventral areas (µm^2^) of each vertebra in methylcellulose-treated wild type and MZ *inppl1a*^*stl*445^ mutants with dashed lines of best fit in black and R^2^ values. Lines of best fit for control wild-type and MZ *inppl1a*^*stl*445^ mutant fish are superimposed from Figure 1 in orange and blue, respectively.

**Figure S4.**
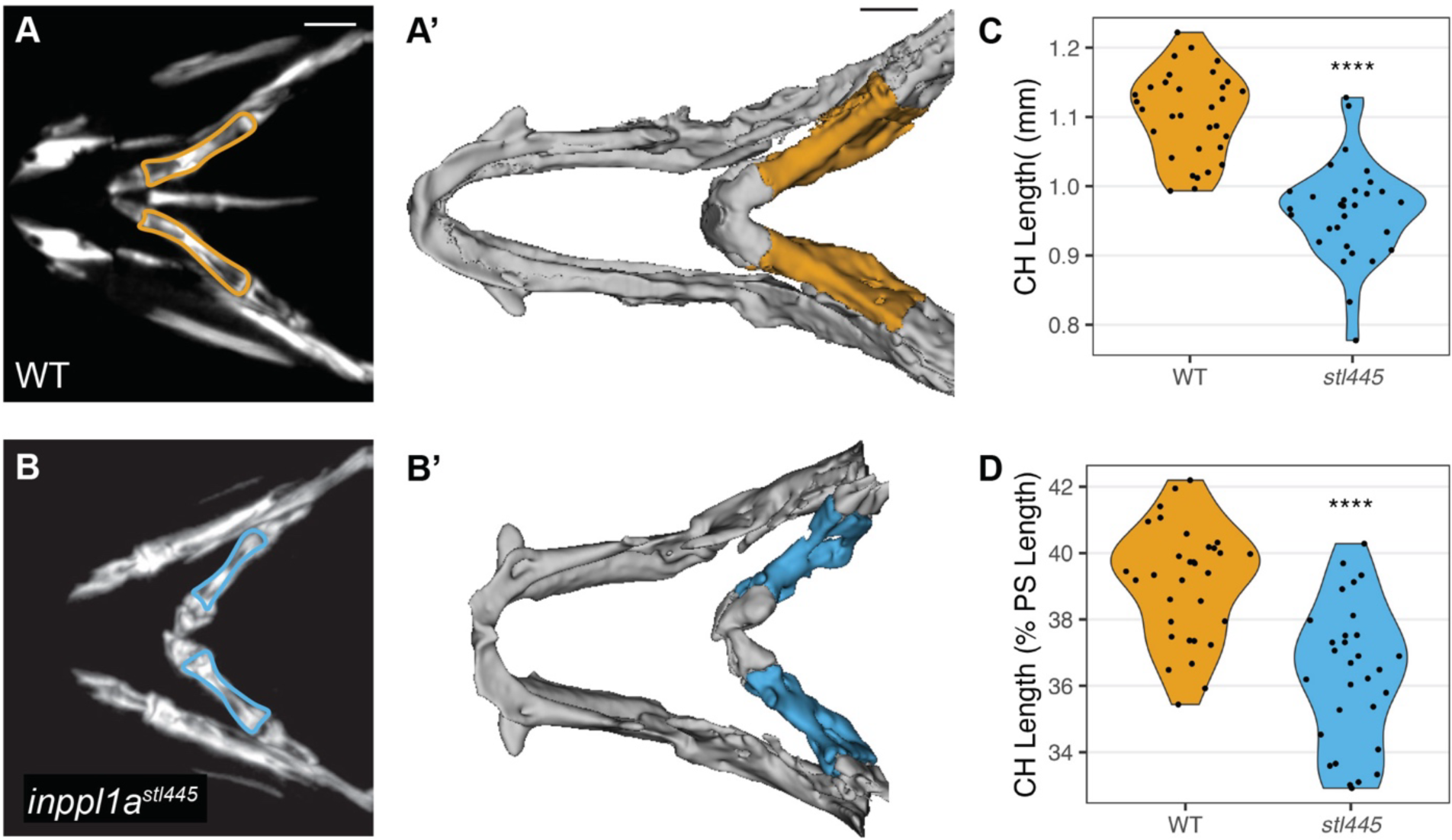
Bone length analysis of the endochondral ceratohyal bones in wild type and MZ inppl1a^stl445^ mutants. **(A-B’)** Representative max projections of MicroCT data of the skull region surrounding the endochondral ceratohyal bones (outlined) in adult wild-type (A) and MZ *inppl1a*^*stl*445^ mutant fish with corresponding 3D renderings (A’-B’). Scale bars = 0.5 mm. **(C)** Quantification of ceratohyal (CH) bone lengths (mm) in wild-type (n = 32, N = 16) and MZ *inppl1a*^*stl*445^ mutant adults (n = 30, N = 15). **** indicates p-value < 0.0001. **(D)** Quantification of ceratohyal (CH) bone lengths normalized as a percent of the anterior parasphenoid (PS) length in individual wild-type (n = 32, N = 16) and MZ *inppl1a*^*stl*445^ mutant (n = 30, N = 15) adults. **** indicates p-value < 0.0001.

